# Large cranial windows distort sensory maps and degrade feature integration in higher-order cortex

**DOI:** 10.64898/2026.02.13.705795

**Authors:** Amber M. Kline, Hiroaki Tsukano, Muneshwar Mehra, Gates P. Schneider, Michellee M. Garcia, Hiroyuki K. Kato

**Affiliations:** Department of Psychiatry, University of North Carolina at Chapel Hill, Chapel Hill, NC 27599, USA; Neuroscience Center, University of North Carolina at Chapel Hill, Chapel Hill, NC 27599, USA; Carolina Institute for Developmental Disabilities, University of North Carolina at Chapel Hill, Chapel Hill, NC 27599, USA

## Abstract

Understanding how sensory information is transformed along cortical hierarchies requires reliable measurement of sensory representations across areas. In vivo two-photon calcium imaging through chronic cranial windows has become an essential approach for characterizing cortical sensory representations with cellular resolution and spatial registration across experiments. However, cortical computations, particularly in higher-order cortices, depend on intricate intracortical circuitry and can be sensitive to subtle tissue perturbations introduced during surgical preparation. Here, we report that implantation of large cranial windows can mechanically compress the intrinsically curved cortical surface, producing systematic distortion that is usually not recognized as overt tissue damage. By comparing sensory maps in the mouse auditory cortex before and after window implantation, we show that larger windows commonly used across laboratories are associated with distorted sensory maps away from the center of the window. Furthermore, both macroscopic and cellular-level imaging reveal a deterioration of feature-selective responses in the secondary auditory cortex (A2) following such distortion. Together, these findings identify an overlooked but controllable surgical factor that can bias measurements of sensory representations and highlight an important consideration for enhancing reproducibility across studies.

## Introduction

In vivo two-photon calcium imaging through chronic cranial windows is now an essential technique in sensory systems neuroscience, enabling simultaneous recording from hundreds to thousands of neurons while preserving their spatial contexts (Holtmaat et al., 2009; Goldey et al., 2014; Kim et al., 2016). With recent advances in wide-field two-photon microscopy, larger cranial windows are increasingly used to image multiple brain regions within the same animal (Sofroniew et al., 2016; Stirman et al., 2016; Rumyantsev et al., 2020; Ota et al., 2021). Combined with genetically encoded calcium indicators (Nakai et al., 2001; Chen et al., 2013; Zhang et al., 2023), these approaches have accelerated efforts to define distinct sensory representations across cortical regions and to probe circuit mechanisms that support area-specific computations.

A key requirement, however, is that the cranial window provides optical access without altering the computations being measured. Because most synaptic input to cortical neurons arises from intracortical circuits, and activity patterns are dominated by recurrent network dynamics (Douglas and Martin, 2004; da Costa and Martin, 2009; Lien and Scanziani, 2013; Kato et al., 2017), even subtle tissue perturbations can alter the intricate circuit and bias measured sensory representations. This concern may be particularly relevant in higher-order cortices, where feature integration depends on distributed interactions across areas. Perturbations that are mild enough to be overlooked but systematic enough to recur within a laboratory could degrade complex feature extraction while leaving basic response properties relatively intact, thereby hampering reproducibility across studies.

These concerns are directly relevant to a discrepancy in reported functional specialization between the primary (A1) and secondary (A2) auditory cortices in mice. We previously found that multi-frequency sounds with coincident onsets preferentially recruit A2 neurons (Kline et al., 2021, 2023). Across intrinsic signal imaging, two-photon calcium imaging, and electrophysiological recordings, A2 responses were strongest for coincident multi-frequency sounds and deteriorated with onset timing shifts as short as 10–15 ms, consistent with coincidence-dependent perceptual integration in human psychophysics (Bregman and Pinker, 1978; Darwin, 1984; Bregman, 1990). In contrast, a recent two-photon imaging study reported a much less pronounced functional difference between A1 and A2 (Chen et al., 2025), raising questions about hierarchical integration of multi-frequency sounds across cortical regions. In addition, in vivo two-photon calcium imaging studies have reported a broad range in the fraction of tone-responsive neurons in A1, from approximately 5% to 40% (Issa et al., 2014; Liu et al., 2019; Aponte et al., 2021). This variability is difficult to account for by differences in calcium indicators or response-detection criteria, suggesting that additional experimental factors may systematically influence the measured cortical sensory representations.

One potential contributor to such differences in findings is variation in surgical preparation. Beyond the well-recognized risks of overt trauma during craniotomy or heat from drilling, subtler mechanical factors may also systematically compromise circuit integrity. In particular, because the lateral cortex is intrinsically curved, fitting a large, flat glass window can impose mechanical compression and perturb neuronal organization and circuit computation. However, despite the widespread use of cranial windows, the extent to which window geometry affects neuronal circuit activity has not been quantified.

To test this directly, we measured auditory cortex tonotopic maps before and after window implantation and related the changes to window size. We show that larger windows distort the sensory map by suppressing responses in the middle of the window and displacing the map away from the center. Using both macroscopic mapping and cellular-resolution imaging, we further find that such distortion is accompanied by a deterioration of A2 neurons’ preference for coincident multi-frequency sounds. Notably, commonly used window designs in auditory cortical imaging fall within the range in which we observe these effects, suggesting that compression-related changes could contribute to systematic alterations in sensory representations across laboratories.

Together, these results identify cranial window size as a controllable source of distortion in sensory maps and degradation of higher-order feature representations. By quantifying how window dimensions relate to map integrity, our study provides a practical framework for minimizing surgical preparation-dependent variability in chronic imaging experiments.

## Results

### Transcranial intrinsic signal imaging reproducibly maps harmonic responses to A2

To quantify distortion of auditory cortical maps following cranial window implantation, we first established baseline maps in intact brains using transcranial intrinsic signal imaging and assessed their reproducibility across animals and experimenters. Imaging was performed in 44 mice by four independent experimenters who were not involved in generating the imaging data from our previous studies, which demonstrated preferential representation of multi-frequency sounds in A2 (Kline et al., 2021, 2023).

Cortical responses were measured as changes in reflectance to red illumination evoked by pure tones (3, 10, and 30 kHz; 75 dB SPL) and by harmonic sounds, including both artificial harmonic stacks (2–40 kHz; fundamental frequency (F_0_) 2 or 4 kHz; 75 dB SPL total) and natural mouse harmonic vocalizations (75 dB SPL total) (Grimsley et al., 2011; Kline et al., 2021) (Fig. 1A, B). Based on pure tone responses, we performed semiautomated segmentation of auditory cortical area boundaries into the primary auditory cortex (A1), ventral auditory field (VAF), anterior auditory field (AAF), and secondary auditory cortex (A2) (Fig. 1C) (Narayanan et al., 2023).

**Figure 1.**
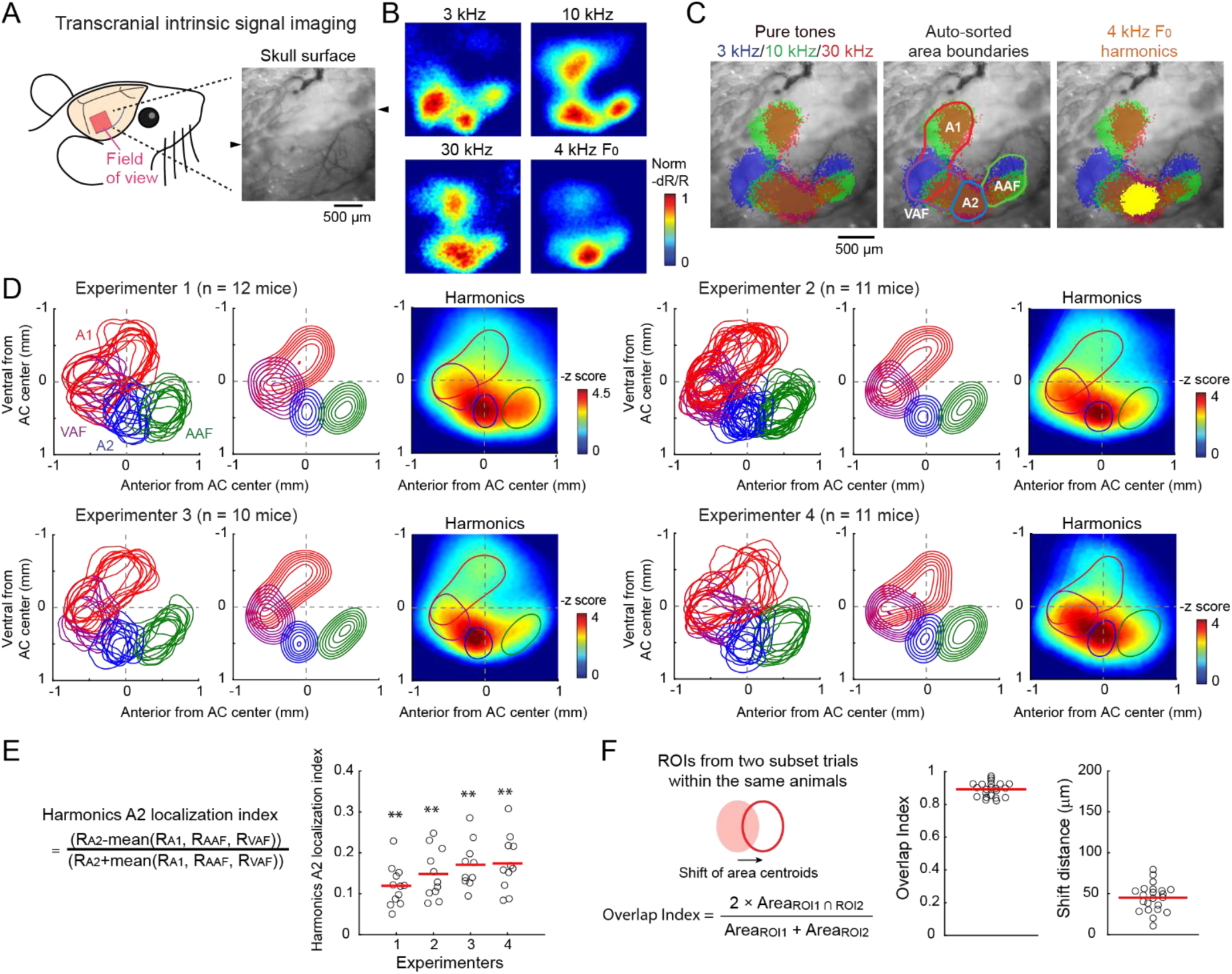
Reproducible mapping of harmonic responses to A2. **(A)** Left, schematic illustrating the field of view used for intrinsic signal imaging. Right, representative image of the skull surface. Black arrowheads indicate a ridge in the skull. **(B)** Intrinsic signal responses to 3, 10, 30 kHz pure tones and 4 kHz-F0 harmonics in the same field of view shown in (A). Heat maps show response amplitudes normalized to the peak. **(C)** Left, thresholded intrinsic signals for pure tone responses superimposed on cortical vasculature imaged through the skull. Middle, auditory cortical area boundaries determined by semiautomated segmentation. A1: primary auditory cortex; AAF: anterior auditory field; VAF: ventral auditory field; A2: secondary auditory cortex. Right, thresholded intrinsic signal responses to 4-kHz F0 harmonics overlaid on pure tone responses. **(D)** Distributions of functionally identified area boundaries and harmonic responses obtained by four independent experimenters (*n* = 12, 11, 10, and 11 mice). Left, cortical area boundaries from all mice superimposed in coordinates around the AC center. Middle, probability distributions of area masks for individual cortical regions. Contours indicate 10% steps, starting from 30%. Right, heat map of harmonic responses averaged across animals, overlaid with the 50% contour of the area probability distribution. **(E)** Scatter plot showing the A2-localization index of harmonic responses for individual experimenters. Red lines indicate the mean. \*\**p* < 0.01 (two-sided Wilcoxon signed-rank test with Bonferroni correction). **(F)** Left, schematic illustrating the comparison of area ROIs derived from two non-overlapping trial subsets within the same animals. Middle, scatter plot showing the Overlap Index between ROI pairs. Right, scatter plot showing absolute area centroid shift distances, averaged across areas for each mouse. *n* = 22 mice with at least eight trials per stimulus.

Although the absolute stereotaxic positions of the auditory cortex vary substantially across animals (Narayanan et al., 2023), our analysis here focuses on the relative locations of individual areas to quantify changes that may be caused by cranial window implantation. We defined the center of the auditory cortex (“AC center”) as the midpoint of the bounding box enclosing all identified auditory cortical areas and expressed the positions of individual areas relative to this reference. Across animals and experimenters, the relative spatial distributions of the four auditory areas, visualized as either overlaid regions of interest (ROIs) or their probability distributions, were highly reproducible and consistent with previous macroscopic imaging studies (Issa et al., 2014; Liu et al., 2019; Aponte et al., 2021; Kline et al., 2021; Narayanan et al., 2023) (Fig. 1D) (except for differences in interpretations regarding the gap between A1 and AAF, which is filled in some studies that assign preferred frequencies to all pixels regardless of response magnitude (Kalatsky et al., 2005; Romero et al., 2020)).

Mapping harmonic responses onto the tonotopically defined cortical map revealed preferential activation of A2, with weaker activity extending into adjacent regions within the A1, VAF, and AAF, consistent with our previous reports (Fig. 1B, C). Harmonic response maps averaged across animals and aligned to the AC center showed robust A2-preferring patterns across experimenters (Fig. 1D, heatmaps). To quantify the A2 preference of harmonic responses, we calculated an A2 localization index defined as (R_A2_ – mean(R_A1_, R_VAF_, R_AAF_))/(R_A2_ + mean(R_A1_, R_VAF_, R_AAF_)), which ranges from +1 (exclusive localization to A2) to -1 (exclusive localization to non-A2 regions). This index was positive in all 44 animals, demonstrating robust and reproducible preferential activation of A2 in intact brains (0.15 ± 0.01, *p* = 7.6 × 10^-9^, two-sided Wilcoxon signed-rank test; for individual experimenters, *p* = 0.0020, 0.0039, 0.0078, 0.0039; two-sided Wilcoxon signed-rank test with Bonferroni corrections) (Fig. 1E). Results obtained from mice on a C57BL/6J background were indistinguishable from those obtained from B6-Cdh23 mice, which do not exhibit age-related hearing loss (Supplementary Fig. 1).

To assess within-animal reproducibility, we analyzed mice with at least eight trials per stimulus by independently generating area ROIs from two non-overlapping subsets of trials. We quantified reproducibility using an Overlap Index, defined as 2 × A_ROI1∩ROI2_ / (A_ROI1_ + A_ROI2_), where A denotes the area of ROIs derived from each trial subset. This index ranges from 0 (no overlap) to 1 (complete overlap). We additionally quantified spatial shifts of area centroids between the two ROI sets. The Overlap Index was consistently high (*n* = 22 mice; 0.89 ± 0.01), and centroid shifts were small (45.1 ± 3.6 μm), which were much smaller than typical area dimensions (Fig. 1F). Therefore, while auditory cortical maps vary widely across animals (Narayanan et al., 2023), the maps remain consistent across measurements within the same animals. Together, these values define an upper bound of map consistency within intact brains and provide a reference for evaluating map distortion following cranial window implantation in subsequent analyses.

### Cranial window implantation distorts the auditory cortical maps away from the center

Having established reproducible auditory cortical mapping and preferential activation of A2 by multi-frequency sounds in intact brains, we next examined how tonotopic maps change following cranial window implantation. We reanalyzed mice from our previous two-photon calcium imaging study (Kline et al., 2021) together with newly collected animals, all of which were used for two-photon calcium imaging via virally transduced GCaMP. In all experiments, transcranial intrinsic signal imaging was performed prior to window implantation to guide viral injections and window placement. A second round of intrinsic signal imaging was conducted through the glass window approximately two weeks after implantation to confirm the map integrity before two-photon imaging. In our earlier work, mice exhibiting substantial post-implantation map distortion were excluded from analysis, as previously described. In the present study, we included these previously rejected mice to evaluate factors influencing tonotopic map preservation in an unbiased manner.

To quantify map alterations across 50 mice, we independently generated area ROIs from intrinsic signals obtained before and after window implantation (Fig. 2A–C). After determining the best translation of the pre-window maps that maximized their overlap with post-window maps (see Materials and Methods), two sets of area ROIs were overlaid to visualize changes. Alignment accuracy was also validated by matching blood vessel patterns (black arrowheads in Fig. 2A–C), although vascular landmarks were less visible in some animals with extensive skull vasculatures (e.g., Mouse 1 in Fig. 2A).

**Figure 2.**
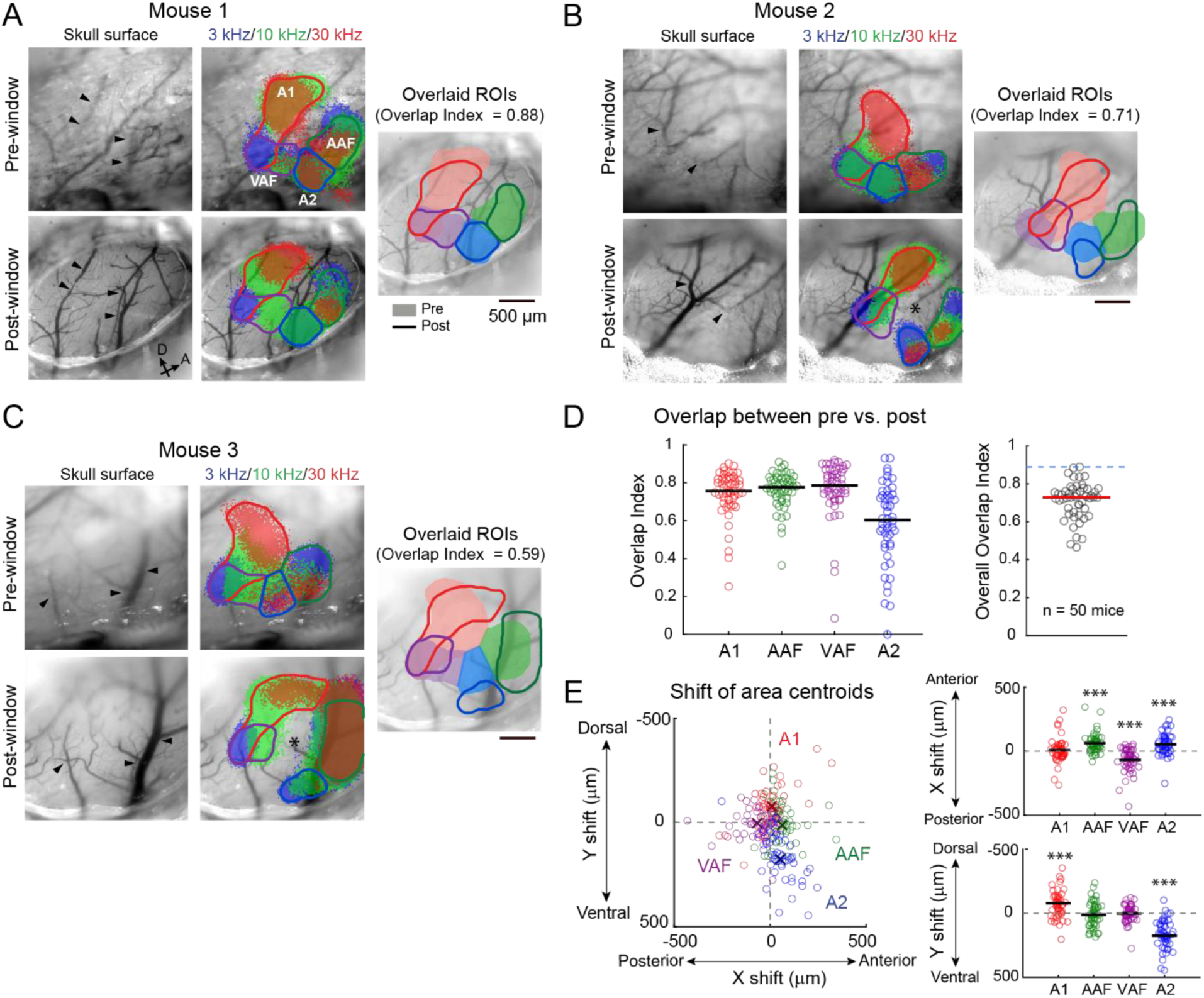
Cranial window implantation distorts tonotopic maps in the auditory cortex. **(A)** Left, images of cortical vasculature acquired before (top) and after (bottom) cranial window implantation in the same mouse. Black arrowheads indicate the same blood vessels identified across imaging sessions. Note that in this animal, most vasculature imaged before window implantation was located within the skull. Middle, intrinsic signal responses to pure tones at 3, 10, and 30 kHz before (top) and after (bottom) window implantation, with overlaid autosorted area boundaries. Right, area boundaries derived from pre-window (shadings) and post-window (lines) imaging superimposed on the post-window vasculature image, illustrating the high overlap between the two sets of area ROIs (Overlap Index = 0.88). **(B)** Same as in (A), but for a mouse exhibiting moderate alteration of tonotopic map organization following window implantation (Overlap Index = 0.71). Asterisk shows a central region with reduced sound responses after window implantation. **(C)** Same as in (A), but for a mouse exhibiting severe alteration of tonotopic map organization following window implantation (Overlap Index = 0.59). **(D)** Left, scatter plot showing the Overlap Index between pre- and post-window area ROIs, calculated separately for individual auditory cortical areas. Black lines indicate means. Right, scatter plot showing the overall Overlap Index averaged across areas for each mouse. The red line indicates the mean. The blue dotted line indicates the upper bound of map reproducibility determined in Fig. 1F. *n* = 50 mice. **(E)** Left, scatter plot showing centroid shifts for individual auditory cortical areas along the anterior-posterior (x-axis) and dorsoventral (y-axis) dimensions. Crosses indicate the mean displacement for individual areas. Top right, centroid shifts along the anteroposterior axis. Bottom right, centroid shifts along the dorsoventral axis. \*\*\**p* < 0.001 (two-sided Wilcoxon signed-rank test with Bonferroni correction).

This analysis revealed substantial variability across animals. In some mice, pre- and post-window maps aligned closely (Mouse 1: Overlap Index = 0.88; Fig. 2A), whereas others showed pronounced distortion (Mouse 2: Overlap Index = 0.71; Mouse 3: Overlap Index = 0.59; Fig. 2B, C). Distorted maps typically exhibited a loss of responses near the map center (asterisks in Fig. 2B, C), accompanied by outward displacement of post-window areal ROIs. We quantified distortions using the Overlap Index and the areal centroid shifts between pre- and post-window maps. When calculated separately for individual areas (Fig. 2D, left) or averaged across areas within each mouse (Fig. 2D, right), Overlap Index values spanned a wide range, indicating the heterogeneity in preservation of tonotopic organization (overall Overlap Index: 0.47–0.89, 0.71 ± 0.01; mice accepted in our paper: 0.62–0.88, 0.74 ± 0.01). A2 tended to exhibit a lower Overlap Index than other areas, likely reflecting its small size and proximity to the map center.

Analysis of area centroid shifts relative to the AC Center revealed a consistent pattern of displacement away from the AC Center. Specifically, A1, VAF, AAF, and A2 centroids were shifted in dorsal, caudal, rostral, and ventral directions, respectively (Fig. 2E). In addition to these distortions away from the center, some animals exhibited asymmetric, side-biased distortions, likely reflecting the relative position of the auditory cortex within the window and the resulting nonuniform compression (Supplementary Fig. 2). Together, these results clearly demonstrate that cranial window implantation can systematically distort auditory cortical maps, particularly away from the map center, motivating us to examine the factors that promote map preservation.

### Cranial window size predicts distortion of auditory cortical maps

Based on the consistent suppression of sound responses around the map center and the outward displacement of auditory cortical areas, we hypothesized that mechanical compression near the window center underlies the observed distortions. Because the brain surface is intrinsically curved, forcing a flat glass necessarily imposes mechanical constraints, with compression expected to be greatest near the center of the window. While the dorsal cortex, where much of wide-field two-photon imaging is conducted, is relatively flat, curvature is more pronounced in the lateral cortex, where the auditory cortex is located. Indeed, fitting a straight 3-mm surface, a window size often used in the mouse auditory cortex, around the auditory cortex in the Allen Common Coordinate Framework (CCF) (Wang et al., 2020) illustrates substantial curvature, particularly along the dorsoventral axis (Fig. 3A, Supplementary Fig. 3). This geometry predicts greater compression along the dorsoventral than the rostrocaudal direction. Accordingly, window shapes with a larger anteroposterior (D_AP_) than dorsoventral diameter (D_DV_) should minimize compression while maximizing imaging area (Fig. 3A, right). In our experiments, we used oval windows with such geometry (*n* = 50 mice, D_AP_ = 2.84 ± 0.04 mm; D_DV_ = 2.00 ± 0.04 mm; mice included in our previous work: *n* = 23 mice, D_AP_ = 2.77 ± 0.06 mm; D_DV_ = 1.85 ± 0.05 mm). Although the resulting window area (4.49 ± 0.14 mm^2^; mice included: 4.05 ± 0.18 mm^2^) was substantially smaller than that of commonly used 3-mm circular windows (7.07 mm^2^), we nevertheless examined whether window dimensions predicted the degree of map distortion.

**Figure 3.**
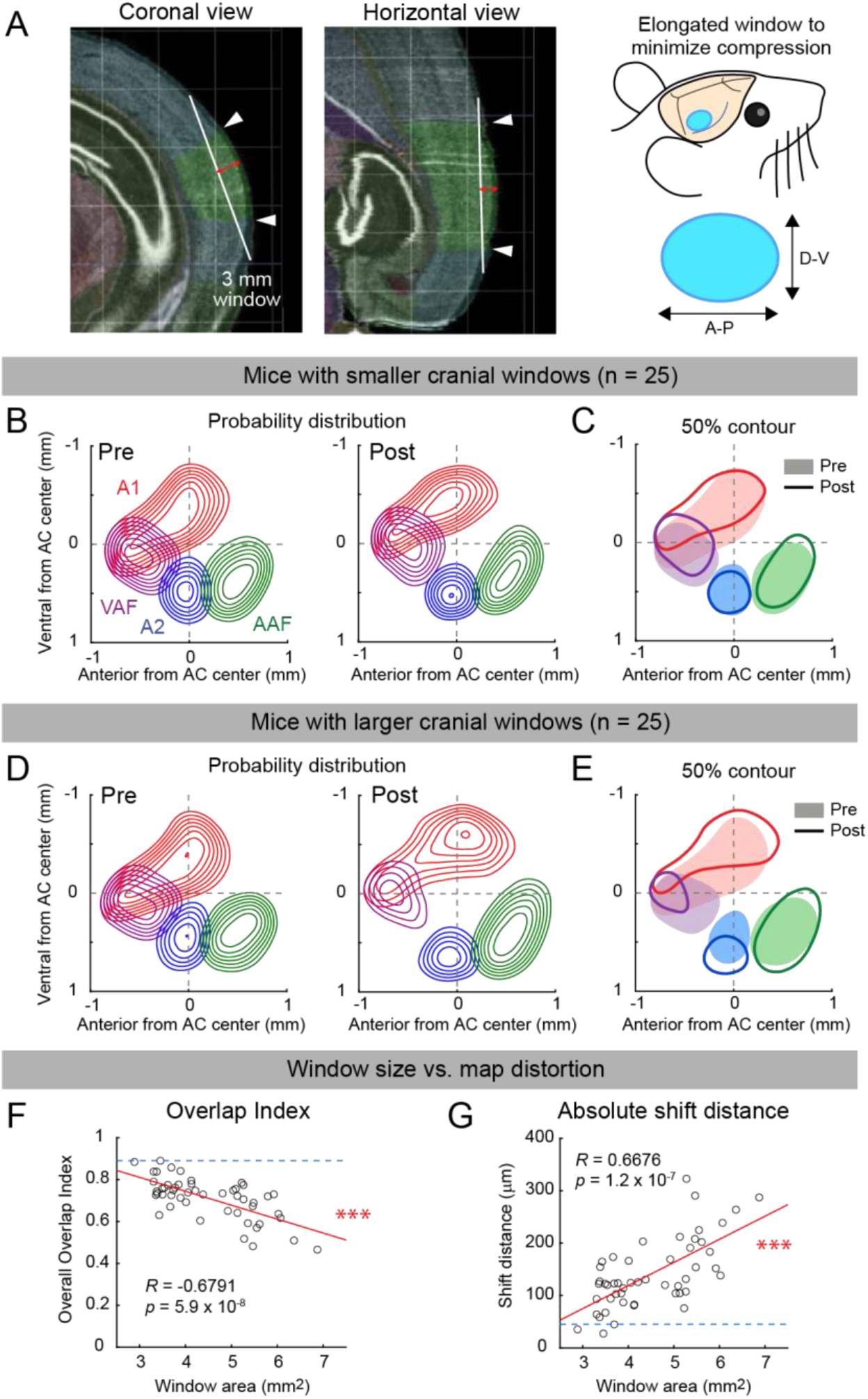
Cranial window size predicts distortion of tonotopic organization. **(A)** Left, cortical structure surrounding the auditory cortex in the Allen CCF, shown in coronal (left) and horizontal (right) views. White lines indicate the diameter of a 3-mm cranial window positioned against the curved cortical surface. Red arrows indicate the resulting compression. White arrowheads and green shading denote the boundaries of the auditory cortex as defined in the Allen CCF. Right, schematics illustrating an oval cranial window elongated along the anteroposterior axis, which is predicted to minimize cortical compression based on the geometry of the Allen CCF. **(B)** Probability distributions of area masks for individual cortical regions around the AC center before (left) and after (right) window implantation in 25 mice with cranial window areas smaller than the median (4.17 mm^2^). Contours indicate 10% steps, starting from 30%. **(C)** 50% contour lines of the area probability distributions for individual auditory cortical regions overlaid between pre-window (shading) and post-window (contours) sessions for the same mice as in (B). **(D, E)** Same as (B, C), but for 25 mice with cranial window area larger than the median. **(F)** Overall Overlap Index plotted as a function of cranial window area for individual mice. Red lines indicate linear regression fits. Blue dotted lines denote the upper bound of map reproducibility determined in Fig. 1F. \*\*\**p* = 5.9 × 10^-8^. **(G)** Centroid shift distances averaged across all auditory cortical regions plotted as a function of cranial window area for individual mice. \*\*\**p* = 1.2 × 10^-7^. *n* = 50 mice.

We first divided the 50 mice into two groups based on window area. In mice with window areas smaller than the median (4.17 mm^2^), probability distributions of cortical areas were largely preserved after window implantation, with only modest outward shifts of the 50% contour lines (Fig. 3B, C). In contrast, mice with window areas larger than the median showed pronounced outward displacement and broadening of areal probability distribution, reflected by lower peak probability and larger shifts of the 50% contour lines (Fig. 3D, E). Note that the 50% contour lines of the probability distribution do not mean the averaged shapes of individual areas; smaller post-window 50% contours reflect greater location variability across animals rather than area shrinkage.

We next directly compared the window area against the measures of map distortion. Window area was strongly correlated with both the Overlap Index (*R* = -0.68, *p* = 5.9 × 10^-8^; Fig. 3F) and the shift distance of areal centroids (*R* = 0.67, *p* = 1.2 × 10^-7^; Fig. 3G). Linear regression indicated that both measures approached the upper bound of map consistency observed in intact brains (Fig. 1F; blue dotted lines in Fig. 3F, G) at window areas of approximately 3-4 mm^2^, while distortion increased monotonically with larger windows. These relationships remained true when the distortion measures were quantified separately for individual areas (Supplementary Fig. 4), with A2 consistently showing the largest susceptibility.

To gain more insights into the optimal window dimensions for mouse auditory cortex imaging, we examined distortion metrics as a function of D_DV_ and D_AP_ independently (Supplementary Fig. 5A, B, D). Both dimensions correlated strongly with distortion measures (Overlap Index vs. D_DV_: *R* = -0.60, *p* = 3.5 × 10^-6^; vs. D_AP_: *R* = -0.66, *p* = 2.3 × 10^-7^; shift distance vs. D_DV_: *R* = 0.60, *p* = 4.9 × 10^-6^; vs. D_AP_: *R* = 0.64, *p* = 5.2 × 10^-7^). Since D_DV_ and D_AP_ covary in our dataset, multiple regression does not disentangle their independent contributions. Instead, we reasoned that the intrinsic geometry around the auditory cortex defines an optimal D_DV_/D_AP_ ratio (r_opt_) that maximizes the window area while minimizing brain compression. In this scenario, compression would be governed by whichever dimension exceeds the optimal geometry—that is, by Max([D_DV_, r_opt_ × D_AP_]) (See Materials and Methods). To determine r_opt_, we varied the ratio r between 0 and 2 and identified the value that maximized the absolute correlation between Max([D_DV_, r × D_AP_]) and each distortion measure. This analysis yielded an identical r_opt_ of 0.75 for both Overlap Index and shift distance (Supplementary Fig. 5C, E, left), indicating that the optimal ratio of D_DV_/D_AP_ for the auditory cortical geometry is 0.75.

Plotting distortion measures against this best-explaining variable, Max([D_DV_, 0.75 × D_AP_]), indicates that window dimensions near D_DV_ = 2 mm minimize cortical map distortion while capturing all auditory cortical areas (Supplementary Fig. 5C, E, right). With D_DV_ fixed at 2 mm, D_AP_ can be extended up to ∼2.67 mm (2/0.75) without causing additional distortion.

Together, these analyses demonstrate that cranial window size and geometry are strong predictors of auditory cortical map distortion, with substantial dislocations occurring at window sizes often used in mouse auditory cortex imaging studies.

### Cranial window size predicts deterioration of harmonic responses in A2

We next examined how cortical map distortion affects sensory feature integration in the higher-order auditory cortex, A2. Overlaying harmonic responses to tonotopic maps in the same example animals shown in Figure 2A-C revealed substantial redistribution of harmonic responses following window implantation (Fig. 4A). In mice with preserved tonotopic maps, preferential activation of A2 by harmonic stimuli remained intact after window implantation (Mouse 1). In contrast, mice with distorted maps showed a redistribution of harmonic responses to adjacent areas, with the peak responses often shifting away from A2 (Mouse 2, 3).

**Figure 4.**
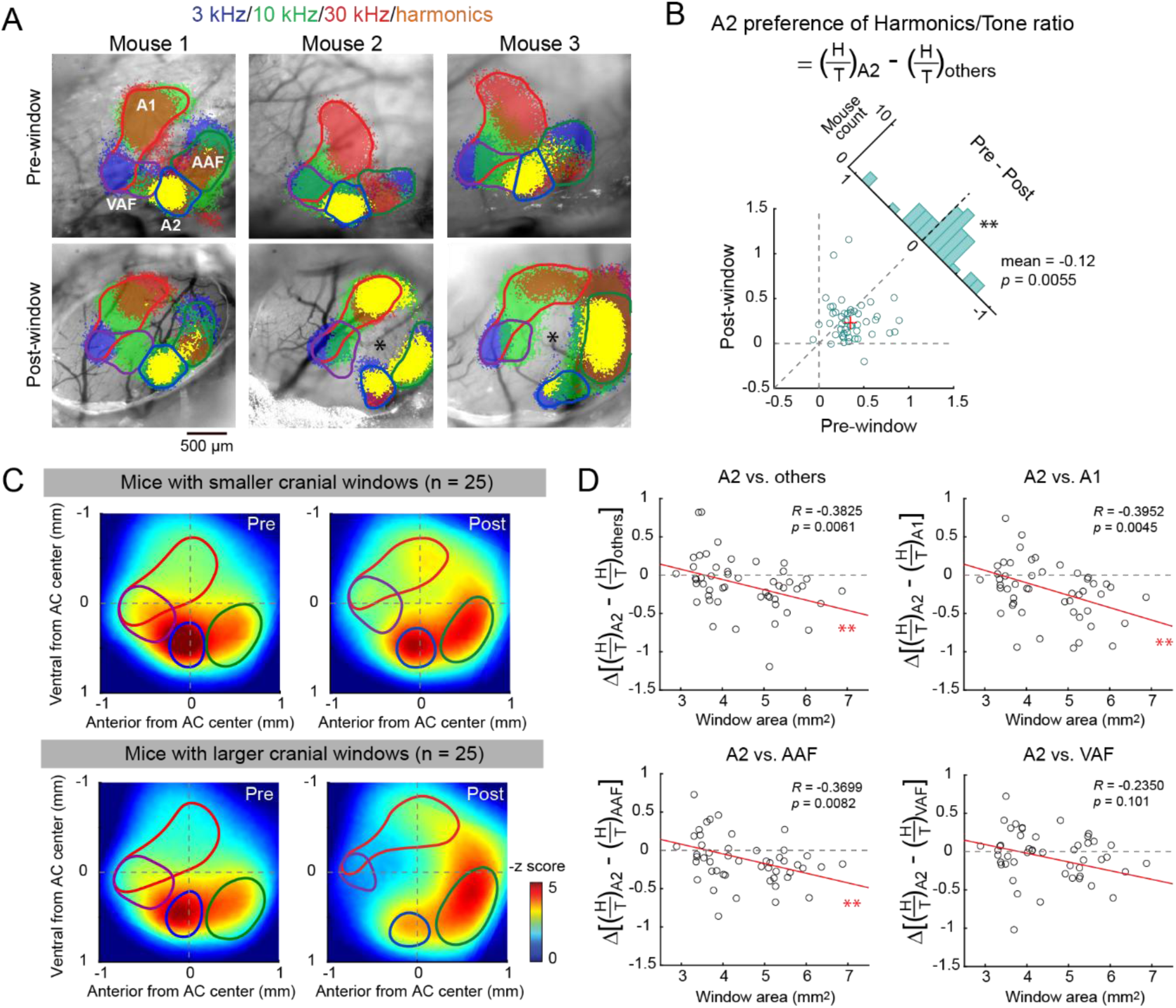
Cranial window size predicts degradation of multi-frequency integration in A2. **(A)** Thresholded intrinsic signal responses to 3, 10, and 30 kHz pure tones and harmonic stimuli, overlaid on cortical vasculature images obtained before (top) and after (bottom) window implantation in the same mice shown in Fig. 2A–C. Lines indicate autosorted area boundaries. Asterisks denote central regions with reduced sound-evoked responses after window implantation. **(B)** Scatter plot showing the A2 preference of the harmonics-to-tone response ratio before and after window implantation. Red cross indicates the mean. The oblique histogram illustrates changes in A2 preference following window implantation. *n* = 50 mice. \*\**p* = 0.0055 (two-sided Wilcoxon signed-rank test). **(C)** Top, heat maps of harmonic responses averaged across 25 mice with cranial window areas smaller than the median (4.17 mm^2^) before (left) and after (right) window implantation, overlaid with 50% contour lines of the area probability distributions for individual auditory cortical regions. Bottom, same as top panels, but for 25 mice with cranial window areas larger than the median. **(D)** Changes in pairwise interareal differences in the harmonics-to-tone response ratio following window implantation, plotted as a function of cranial window area for individual mice. Red lines indicate linear regression fits. Top left, A2 vs. others; top right, A2 vs. A1; bottom left, A2 vs. AAF; bottom right, A2 vs. VAF. One outlier datapoint in A2 vs. VAF (x = 5.1, y = -2.5) lies outside the displayed range. \*\**p* < 0.01.

As absolute response magnitudes are not directly comparable between pre-window transcranial imaging and post-window imaging through glass, we quantified harmonic selectivity using the harmonics-to-tone ratio (H/T). We then defined a harmonic A2 preference index as (H/T)_A2_ – (H/T)_others_ (see Materials and Methods). Before window implantation, this index was consistently positive across animals, indicating preferential harmonic activation of A2 in intact brains (*n* = 50 mice; Pre-window axis: 0.35 ± 0.03, *p* = 9.1 × 10^-10^; two-sided Wilcoxon signed-rank test) (Fig. 4B). After window implantation, the same mice showed a significant reduction in this index to 0.23 ± 0.04 (pre vs. post: *p* = 0.0055), indicating deterioration of area-selective harmonic integration.

To assess the dependence of this effect on window size, we again divided mice into two groups based on window area and mapped harmonic responses onto tonotopically defined cortical areas before and after window implantation (Fig. 4C). In mice with window areas smaller than the median (4.17 mm^2^), harmonic response peaks remained localized within A2, despite modest redistribution across areas. In contrast, mice with larger windows showed a marked reduction of harmonic responses in A2, with response peaks shifting toward AAF. This apparent shift toward AAF may be influenced by the positioning of the auditory cortex in our preparation, which was biased rostrally within the window to place A1 near the window center, resulting in relatively milder tissue compression in AAF.

We next directly compared the window area against the changes in harmonic responses localization to A2. Linear regression revealed a significant negative correlation between window area and the change in harmonic A2 localization index after window implantation (Δ[(H/T)_A2_-(H/T)_others_]_post-pre_), indicating that the A2-localization of harmonic responses deteriorated with the increased window size (*R* = -0.3825, *p* = 0.0061; Fig. 4D, top-left). This effect was not driven solely by increased harmonic responses in AAF; similar window size dependence was observed when localization indices were computed for A2 relative to A1, AAF, or VAF individually (A2-A1: *R* = -0.40, *p* = 0.0045; A2-AAF: *R* = -0.37, *p* = 0.0082; A2-VAF: *R* = -0.24, *p* = 0.101).

As observed for tonotopic map distortion (Fig. 3), the change in A2 harmonic response localization (post - pre) approached zero at window areas of approximately 3-4 mm^2^, indicating minimal deterioration at these sizes. Together, these results demonstrate that implantation of large cranial windows not only distorts tonotopic organization but also deteriorates sensory feature integration in A2.

### Cranial window size predicts deterioration of harmonic integration at the cellular level

We next asked whether the macroscopic deterioration of cortical harmonic responses is also reflected at the cellular level. We reanalyzed in vivo two-photon calcium imaging data from mice presented with multi-frequency harmonic stacks while imaging layer 2/3 pyramidal neurons (*n* = 18 mice; A1 and A2 were imaged in 16 and 15 mice, respectively). Excitatory pyramidal neurons were identified as tdTom-negative neurons in VGAT-Cre × Ai9 mice. Artificial harmonic stacks (F_0_ = 2, 2.8, 4, 5.7, or 8 kHz; harmonics up to 40 kHz) were presented at a total intensity of 70 dB SPL. To probe coincidence-dependent sound integration, onset timing shifts were introduced between the lower- and upper halves of their frequency components (Δonset: timing of lower compared to upper half, -45 to +45 ms in 15-ms steps) (Fig. 5A) (Kline et al., 2021).

**Figure 5.**
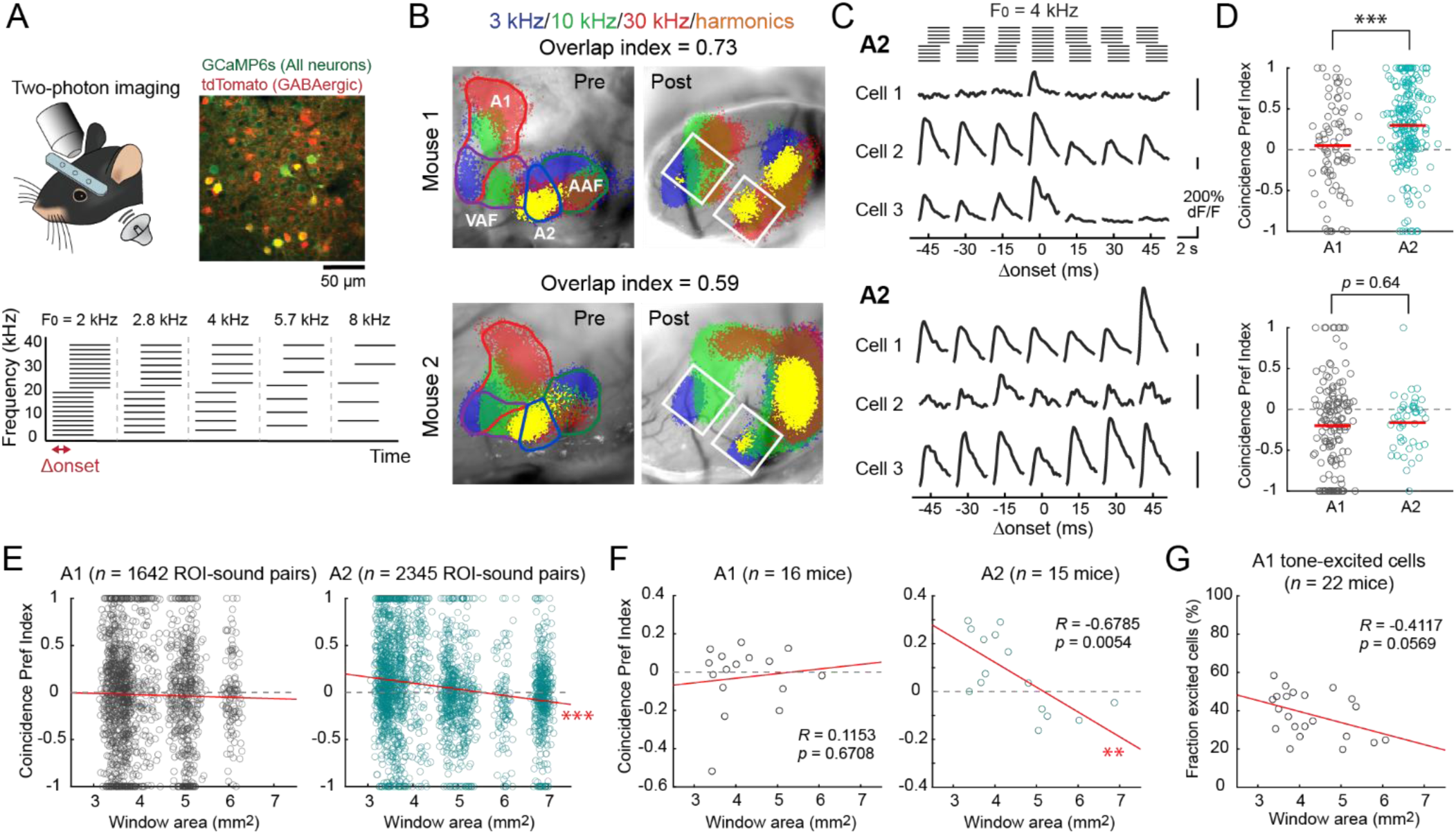
Cranial window size predicts deterioration of harmonic integration at the cellular level. **(A)** Top, representative in vivo two-photon image of layer 2/3 neurons in A2. Bottom, schematic illustrating artificial harmonic stimuli with different F0 and Δonset. **(B)** Intrinsic signal responses to pure tones and harmonic stimuli before (left) and after (right) window implantation in two representative mice, showing better (top) and worse (bottom) preservation of A2 harmonic responses. In Mouse 1, part of A1 lies outside the cranial window. White squares indicate the fields of view used for two-photon calcium imaging. **(C)** Responses of representative layer 2/3 pyramidal neurons in A2 to harmonic stimuli with different Δonsets in the same mice shown in (B). **(D)** Coincidence Preference Index for all cell-stimulus pairs in A1 and A2 from the same mice as in (B, C). A significant difference in Coincidence Preference Index between A1 and A2 was observed only in the mouse with preserved A2 harmonic responses. Mouse 1: \*\*\**p* = 2.87 × 10^-4^; Mouse 2: *p* = 0.64 (two-sided Wilcoxon rank-sum test). **(E)** Coincidence Preference Index from all mice plotted as a function of cranial window size for A1 (left) and A2 (right). Red lines indicate linear regression fits. A1: *n* = 1,642 responsive ROI-sound pairs, *p* = 0.40; A2: *n* = 2,345 responsive ROI-sound pairs, \*\*\**p* = 1.2 × 10^-17^. **(F)** Coincidence Preference Index averaged within individual mice plotted as a function of cranial window size for A1 (left) and A2 (right). A1: *n* = 16 mice, *p* = 0.67; A2: *n* = 15 mice, \*\**p* = 0.0054. **(G)** Fraction of pure tone-responsive neurons in A1 plotted as a function of cranial window size. *n* = 22 mice, *p* = 0.057.

For individual neurons, we quantified coincidence dependence by calculating the Coincidence Preference Index, defined as (C - S)/(C + S), where C and S represent the response amplitude triggered by coincident and temporally shifted sounds, respectively (see Materials and Methods). In our previous study, this metric revealed sharply tuned responses of A2, but not A1, neurons to coincident sounds, which deteriorated with Δonsets as short as 15 ms, mirroring coincidence-dependent perceptual binding in human psychophysics experiments. Consistent with these findings, a representative mouse with a small cranial window and preserved harmonic responses in A2 at the macroscopic level (Fig. 5B, Mouse 1) exhibited strong coincidence preference in A2 even at the cellular level. In this animal, the Coincidence Preference Index was significantly higher in A2 than in A1 (Fig. 5C, D; *p* = 2.9 × 10^-4^, two-sided Wilcoxon rank-sum test). In contrast, another mouse with marked deterioration of harmonic responses at the macroscopic level (Mouse 2) showed no difference in coincidence preference between A1 and A2 (*p* = 0.64), indicating a loss of area-specific temporal integration.

We next examined how coincidence preference in individual neurons depended on cranial window size across mice. In A2, the Coincidence Preference Index showed a significant negative correlation with window area, whereas no such relationship was observed in A1 (A1: *R* = -0.0212, *p* = 0.40; A2: *R* = -0.1764, *p* = 1.2 × 10^-17^) (Fig. 5E). While there is substantial heterogeneity across individual neurons, population-level differences between A2 and A1 were evident at window areas of 3–4 mm^2^. In contrast, at larger window sizes (around 6–7 mm^2^), coincidence preference was indistinguishable between the two areas. To account for within-animal clustering, we repeated the analysis by averaging the indices across neurons within each animal and found consistent results (A1: *R* = 0.12, *p* = 0.67; A2: *R* = -0.68, *p* = 0.0054) (Fig. 5F).

Finally, we asked whether cranial window size could also contribute to variability in the fraction of tone-responsive neurons reported across in vivo two-photon GCaMP calcium imaging studies, which ranges from ∼5% (Liu et al., 2019) to 30–40% (Issa et al., 2014; Aponte et al., 2021). We quantified the fraction of significantly tone-excited layer 2/3 pyramidal neurons in A1 across 22 mice. Responses were measured using 17 pure tone frequencies (4–64 kHz, log-spaced) and three sound levels (30, 50, and 70 dB SPL). Although not statistically significant, the fraction of excited neurons showed a negative trend with increasing window area (*R* = -0.41, *p* = 0.0569) (Fig. 5G). This reduced responsiveness with larger cranial windows may contribute to the low fraction of responsive neurons reported in the previous study using 3-mm diameter circular windows.

## Discussion

With the increasing adoption of wide-field two-photon microscopy, experiments are trending toward larger cranial windows to access multiple cortical regions simultaneously. However, the assumption that “bigger is better” for optical access must be balanced against preservation of native cortical geometry, as forcing a flat glass window against the intrinsically curved brain surface can impose mechanical constraints and alter circuit function. In this study, we identify cranial window size as a crucial and underappreciated determinant of sensory map integrity following window implantation in the mouse auditory cortex. By directly comparing auditory tonotopic maps before and after cranial window implantation, we demonstrate that window implantation can distort tonotopic organization, most notably through the suppression of responses near the window center and the outward displacement of areal locations. The severity of these distortions scaled with cranial window dimensions. Importantly, we further demonstrate that these structural distortions are accompanied by a loss of preferential integration of coincident multi-frequency sounds in A2, evident in both macroscopic imaging and cellular-level responses measured with two-photon calcium imaging. Together, these findings indicate that the effects of large window implantation are not limited to a simple downscaling of response amplitudes or remapping of cortical coordinates, but are also associated with altered neuronal tuning and impaired integration of complex sensory features in higher-order cortex.

### Systematic distortion of cortical maps following large cranial window implantation

We found non-random distortions of cortical maps following implantation of large cranial windows. Across animals, sensory responses were often suppressed near the map center, and the centroids of individual areas were displaced away from the center. This structured displacement is consistent with a mechanical constraint imposed by fitting a flat glass window onto the intrinsically curved cortical surface, where compression is expected to be greatest near the center. Generally, protocols for cranial window implantation recommend applying gentle but sufficient pressure to prevent dural thickening, skull regrowth, or brain motion, while avoiding visible disruption of blood flow. In our preparations, we did not observe overt blockage of surface vasculature during window implantation, suggesting that even pressure levels that do not visibly impede surface blood flow can nonetheless be sufficient to distort functional organization. The biological substrates linking compression to altered maps remain to be determined and could include subtle changes in microvasculature, mechanically induced changes in neuronal intrinsic properties, alterations in local circuit connectivity, or eventual neuronal loss.

Importantly, the observed cortical map distortions were not a simple reduction in response magnitudes. Instead, they were accompanied by degraded area-specific computations in A2, particularly for the integration of coincident multi-frequency sounds. We consider two non-mutually exclusive mechanisms for such deterioration. First, attenuation of pure tone responses in A1, as reflected by a reduced fraction of tone-responsive neurons, may alter the transmission of component tone information that is required for downstream multi-frequency integrators. Second, due to the displacement of the functional A2 location, the detected new A2 location may not possess the A2-intrinsic circuit wiring, including lateral and inter-areal interactions, that support higher-order feature integration.

Finally, we note that the reduced responsiveness near the center of compression may not be readily detectable in best frequency maps that assign a preferred sensory stimulus to each pixel regardless of response magnitude (Kalatsky et al., 2005; Romero et al., 2020). While these approaches provide fine-grained tuning information, we suggest that they be accompanied by magnitude maps to visualize low-responsive regions that may reflect genuine circuit organization (e.g., the low-responsive “gap/center” region between A1 and AAF reported in magnitude-based maps (Issa et al., 2014; Tsukano et al., 2015; Liu et al., 2019; Narayanan et al., 2023)) or arise from surgical perturbations.

### Methodological factors affecting multi-frequency integration in A2

Our study was motivated by differences between our previous reports (Kline et al., 2021, 2023) and a recent study by Chen et al., which reported less pronounced functional differences between A1 and A2 (Chen et al., 2025). Notably, although Chen et al. found a significantly higher fraction of harmonic stack-responsive neurons in A2 L2/3 than in A1 L2/3 or L4, they concluded that harmonic-sensitive neurons are equally distributed across regions and layers. While differences in stimulus design (e.g., their 1-s harmonic stacks may reduce the contribution of onset coincidence compared to our 100–200 ms stimuli) or housing environments could contribute, our current study points to an additional, potentially important factor: cranial window-induced cortical compression, which depends on window size and geometry.

Across animals, both tonotopic map integrity and A2’s preferential responses to coincident multi-frequency sounds deteriorated systematically with increasing window size. In our previous studies, we routinely compared tonotopic maps before and after window implantation and excluded mice with substantial map distortion; consequently, the included animals had average window dimensions of 1.85 mm (DV) × 2.77 mm (AP) (area ∼4.01 mm²). By comparison, Chen et al. used a 3-mm circular window (7.07 mm^2^) with a larger 4-mm craniotomy, which could plausibly result in greater cortical compression. Although the effective degree of compression depends on surgical details (see below), our measurements suggest that window sizes in this range are associated with pronounced map distortion (Fig. 3F, G), reduced multi-frequency integration in A2 (Figs. 4–5), and attenuated tone responses in A1 (Fig. 5G). Thus, differences in window size and the resulting cortical deformation represent one possible contributor to differences in measured areal specialization across studies.

Consistent with this interpretation, prior in vivo two-photon calcium imaging studies have reported substantial variability in the fraction of tone-excited neurons, ranging from 30-40% (Issa et al., 2014; Aponte et al., 2021; this study) to ∼5% in a study from the same group (Liu et al., 2019). Multiple factors may influence this metric, including GCaMP expression strategies, imaging signal-to-noise ratio, and housing environment (however, differences in detection threshold are unlikely to account for this variability; see Materials and Methods). Our data suggest that attenuation of cortical responsiveness associated with large cranial windows could represent an additional contributing factor.

Beyond its direct effects on sensory feature integration, sparsening of sound responses can complicate the interpretation of nonlinearity measures. For example, “harmonic-sensitive neurons,” defined as neurons responsive to harmonics but not to pure tones (Chen et al., 2025), may appear more prevalent when overall tone responsiveness is low. This concern is particularly important given that, across sensory modalities, many studies have shown that the hierarchical distinction between primary and higher-order cortices is graded rather than absolute; “higher-order-like” responses are consistently observed in primary cortices, and vice versa (Condylis et al., 2020; Kline et al., 2021, 2023; Siegle et al., 2021; D’Souza et al., 2022). Preserving robust sensory responsiveness is therefore critical for interpretable comparisons of these graded differences in nonlinear integration across neurons and cortical areas.

### Considerations for minimizing window implantation-related artifacts

Reproducibility is a foundational requirement for cumulative progress in science. Recent efforts to develop standardized experimental paradigms and open-source tools for data acquisition and analysis represent important steps toward this goal (Pachitariu et al., 2016, 2017; Aguillon-Rodriguez et al., 2021). In parallel, it is equally essential to identify and control experimental factors that can systematically bias measurements and data interpretation. While variability can arise from differences in sensory stimuli, brain state, housing environment, or analysis pipelines, our findings indicate that surgical preparation is an additional factor that can directly alter both the physiological properties of individual neurons and computations at the circuit level. Because window implantation practices are typically consistent within a given experimenter or laboratory, they may create a false sense of robustness, as results can replicate internally while differing systematically across groups. Based on our findings, and informed by our own ongoing efforts to optimize surgical preparations, we propose several practical considerations.

First, cranial window dimensions and geometry should be treated as experimental variables rather than neutral technical details. Window size, shape (e.g., circular vs. oval), and orientation can all influence cortical compression, sensory map distortion, and degradation of sensory feature integration. For each brain region of interest, community standards may benefit from considering window dimensions that best match local intrinsic geometry. For the mouse auditory cortex, and likely other regions on the lateral cortical surface, oval windows with a larger anteroposterior than dorsoventral extent are predicted to minimize compression while maximizing coverage. Our analysis identifies an optimal D_DV_/D_AP_ ratio of approximately 0.75, corresponding to window dimensions near 2 mm × 2.7 mm, as a reasonable starting point. Similar geometric considerations likely apply to other regions with strong surface curvature, such as secondary somatosensory cortex, lateral secondary visual cortex, and insular cortex. The dorsal cortex is expected to tolerate larger cranial windows due to its relatively flat curvature. Nevertheless, even in the dorsal cortex, in preparations using extremely large windows such as the whole-dorsal “crystal skull” approaches, care should be taken to ensure that window geometry matches the underlying cortical curvature as closely as possible.

Second, functional quality control measures should be incorporated before committing to cellular-level imaging. When feasible, performing functional cortical mapping before and after window implantation provides a direct means to confirm the preservation of map integrity. Alternatively, assessing basic response properties—such as the fraction of neurons responsive to simple stimuli (e.g., pure tones, orientation gratings, whisker deflections)—and comparing them with established field standards, often available from electrophysiology or prior imaging studies, can provide an indirect but informative check. This quality control is particularly important, as the degree of cortical compression can vary even for identical window sizes, depending on surgical procedures. Over the auditory cortex, the skull contains a ridge at the transition between the dorsal and lateral planes (black arrowheads in Fig. 1A). The extent to which this ridge is flattened, as well as the location of the auditory cortex within the window, can influence local pressure and deformation. Consequently, while our findings are qualitatively robust, the magnitude of cortical distortion may differ across laboratories. Performing functional cortical mapping both before and after window implantation would help identify laboratory-specific conditions that preserve cortical organization.

Finally, we note that physical agitation through the intact skull can distort cortical sensory representations even before craniotomy. In transcranial intrinsic signal imaging, experimenters early in their surgical training sometimes observe distorted tonotopic maps, with harmonic responses spreading away from A2. In our experience, these distortions usually arise from excessive mechanical agitation during skull cleaning, such as applying excessive pressure while wiping with cotton swabs or aggressively scraping connective tissue. While systematic quantification of this effect is challenging, careful reduction of mechanical agitation during these preparatory steps reduces the likelihood of such distortions. Although damage to the brain through the intact skull is rarely considered a critical factor, this observation highlights the sensitivity of cortical tissue to physical perturbations and further underscores the importance of careful surgical handling and functional quality control.

In summary, our results identify a controllable surgical factor that can systematically bias measurements of sensory representations and contribute to cross-laboratory variability. The recommendations outlined here are intended to enhance reproducibility and help distinguish preparation-dependent variability from genuine biological differences. In the future, the development and broader adoption of curved glass windows or soft, silicone elastomer-based windows may provide promising alternatives for minimizing cortical compression in studies of inter-areal interactions across extended cortical regions.

## Materials and Methods Animals

The mouse strains used in this study included: C57BL/6J (JAX 000664), B6N-Cdh^23tm2.1Kjn/Kjn^ (JAX 018399), Slc32a1^tm2(cre)Lowl^/J (VGAT-Cre; JAX 028862), Pvalb^tm1(cre)Arbr^/J (PV-Cre; JAX 017320), Sst^tm2.1(cre)Zjh^/J (Sst-Cre; JAX 013044), Vip^tm1(cre)Zjh^/J (VIP-Cre; JAX 010908), Scnn1a-Tg3-Cre (JAX 009613), Calb2^tm1(cre)Zjh^/J (Calb2-Cre; JAX 010774), Pvalb^tm4.1(flpo)Hze^/J (PV-Flp; JAX 022730), and Gt(ROSA)26Sor^tm9(CAG-tdTomato)Hze^/J (Ai9; JAX 007909). Mice were 6–13 weeks old for those on a C57BL/6J background and 6–20 weeks old for those on a B6-Cdh23 background at the time of experiments. Some animals used in Figures 2-5 were reanalyzed from our previous study (Kline et al., 2021). Both female and male mice were used. Animals were housed at 21°C and 40% humidity, with ad libitum access to food and water. Mice were kept under a reverse light cycle (12–12 h), and all experiments were conducted during the dark cycle. All experimental procedures were approved and conducted in accordance with the Institutional Animal Care and Use Committee at the University of North Carolina at Chapel Hill and the guidelines of the National Institutes of Health.

### Auditory stimuli

Auditory stimuli were generated in Matlab (MathWorks) at a sampling rate of 192 kHz and delivered via a free-field electrostatic speaker (ES1; Tucker-Davis Technologies). Speakers were calibrated over a frequency range of 2–64 kHz to achieve a flat response (±1 dB). Sounds were delivered to the ear contralateral to the imaging site. Stimulus presentation was controlled by Bpod (Sanworks) running on Matlab. All pure tones, including those used as pure tone stimuli and as components of multi-frequency stacks, had 5-ms linear rise and fall ramps at onsets and offsets.

For areal mapping with intrinsic signal imaging, pure tones at 3, 10, and 30 kHz (75 dB SPL, 1-s duration) were presented. In some animals, 60 kHz tones were also tested, and their responses largely overlapped with those to 30 kHz tones. Artificial harmonic stimuli with F_0_ = 2, 4, and 8 kHz were generated using harmonic components between 2–40 kHz. The intensity of each frequency component was equalized and adjusted such that the total intensity, summed in sine phases, was 75 dB SPL. As a natural harmonic stimulus, a recorded mouse harmonic vocalization (“P100_09”) from a published library (Grimsley et al., 2011) was presented at 75 dB SPL. Stimuli were presented at 30-second interstimulus intervals for 5–20 trials.

For two-photon calcium imaging experiments, pure tone pips (1-s duration) covering 17 frequencies (4–64 kHz, log-spaced) were presented at 30, 50, and 70 dB SPL. Artificial harmonic stimuli with F_0_ = 2, 2.8, 4, 5.7, and 8 kHz were generated using harmonic components between 2–40 kHz, with a total intensity of 70 dB SPL and a duration of 100 ms. Some experiments used spectrally jittered harmonics, which were generated by adding random frequency jitter (drawn from a uniform distribution between -0.5 × F_0_ and +0.5 × F_0_) to each component of a ten-tone harmonic stimulus with F_0_ = 4 kHz. Temporally shifted harmonics were generated by shifting the onset timing between the lower- and upper halves of frequency components by -45 to +45 ms in 15-ms steps. Sounds were presented in randomized order at 5-second interstimulus intervals, with each stimulus repeated for five trials.

### Intrinsic signal imaging

Intrinsic signal imaging was conducted to locate the auditory cortical areas, following a previously published protocol (Narayanan et al., 2023). Signals were acquired using a custom tandem lens macroscope consisting of a Nikkor 135 mm 1:2.8 lens paired with either a Nikkor 35 mm or a Nikkor 50 mm 1:1.4 lens, attached to a 12-bit CCD camera (DS-1A-01M30, Dalsa) placed in a sound isolation chamber (Gretch-Ken Industries). Mice were anesthetized with isoflurane (0.8–2%) vaporized in oxygen (1 L/min) and maintained on a heating pad at 36-37°C. The muscle overlying the right auditory cortex was removed, and a custom stainless-steel head bar was secured to the skull using dental cement. For initial mapping prior to craniotomy, the cortical surface was imaged through the intact skull, which was kept transparent with phosphate-buffered saline (PBS). For re-mapping before two-photon calcium imaging, the brain surface was imaged through the implanted glass window. Mice received a subcutaneous injection of chlorprothixene (1.5 mg/kg) prior to imaging. Images of surface vasculature were acquired under green LED illumination (530 nm), and intrinsic signals were recorded at 16 Hz under red illumination (625 nm). Images of reflectance were acquired at a resolution of either 717 × 717 pixels (2.3 × 2.3 mm; 135 + 35 mm lens pair) or 1024 × 1024 pixels (4.4 × 4.4 mm; 135 + 50 mm lens pair). Images collected during the response window (0.5–2 seconds after sound onset) were averaged across trials and normalized to the average image during the pre-stimulus baseline period. Images were Gaussian filtered (σ = 2 pixels) and averaged across trials for each stimulus using the IO and VSD Signal Processor Plugin on Fiji software (https://imagej.net/software/fiji/; https://murphylab.med.ubc.ca/io-and-vsd-signal-processor/) (Harrison et al., 2009). The resulting images were deblurred using a 2-D Gaussian window (σ = 200 μm, corresponding to 68 pixels) using the Lucy-Richardson deconvolution method (Orbach and Cohen, 1983; Issa et al., 2014; Romero et al., 2020) to generate a trial-averaged response intensity map. Individual auditory cortical areas, including A1, AAF, VAF, and A2, were identified based on their characteristic tonotopic organization. Specifically, A1 was identified as the most caudal area whose tonotopic gradient traveled rostrodorsally (low→high), and this area likely includes the ultrasound field (UF) in earlier studies (Stiebler et al., 1997). VAF was identified as the most caudal area with a rostroventrally oriented tonotopic gradient. A1 and VAF converged at their low-frequency poles in most animals (Issa et al., 2014; Liu et al., 2019; Romero et al., 2020; Narayanan et al., 2023). AAF was identified as the most rostral area whose tonotopic gradient traveled caudally, with most mice showing a caudoventral gradient. Finally, A2 was identified as the tone-responsive region located between VAF and AAF, which usually exhibits a weak tonotopic gradient traveling ventrally. For visualization of functional maps in figures, signals were thresholded independently for each stimulus.

## Two-photon calcium imaging

Following transcranial mapping of auditory cortical areas using intrinsic signal imaging, a craniotomy (approximately 2 × 3 mm) was performed over the right auditory cortex, with the dura left intact. Drilling was paused every 1–2 s, and the skull was cooled with PBS to prevent damage from overheating. Virus was injected at 5–10 sites per animal (250 μm deep below the pial surface, 30 nL/site at 10 nL/min). For pyramidal neuron imaging, AAV9-syn-GCaMP6s (2 × 10^12^ genome copies per mL) was injected in VGAT-Cre×Ai9 mice. For other cell type-specific imaging, AAV9-syn-Flex-GCaMP6s (2–4 × 10^12^ genome copies per mL) was injected into PV-Cre, Sst-Cre, or Scnn1a-Cre mice. Although all injected animals were included in pre- and post-window comparisons of intrinsic signals, only pyramidal neuron data were used for cellular-level analyses of sound responses in two-photon calcium imaging. A glass cranial window was placed over the craniotomy and secured with 4% agarose and dental cement. The window assembly consisted of two layers of smaller oval #1 coverslips positioned at the center of a larger oval #1 coverslip. The edges of the larger coverslip rested on the skull, whereas the smaller inner coverslips directly contacted the cortical surface; therefore, we refer to the smaller inner coverslips as the cranial window and used their dimensions for window size quantification. Dexamethasone (2 mg/kg) was administered prior to the craniotomy, and Enrofloxacin (10 mg/kg) and Meloxicam (5 mg/kg) were administered before the mice were returned to their home cages. Two-photon calcium imaging was performed 2–3 weeks after cranial window implantation to ensure an appropriate level of GCaMP6s expression. A second intrinsic signal imaging through the chronic window was conducted 1–3 days before two-photon calcium imaging to examine preservation of tonotopic organization. During calcium imaging, awake mice were head-fixed under a two-photon microscope in a custom-built sound-attenuation chamber. GCaMP6s and tdTomato were excited at 925 nm (InSight DS+, Newport). Images (512 × 512 pixels covering 620 × 620 μm) were acquired with a commercial two-photon microscope (MOM scope, Sutter) running ScanImage software (MBF Bioscience) using a 16× objective (Nikon) at 30 Hz. Imaging was performed in cortical layer 2/3 (150-300 μm below the pial surface). Timings of sound delivery were aligned to imaging frames by recording timing TTL signals in WaveSurfer software (Vidrio). An example field of view in Fig. 5A was generated by overlaying signals from two channels using Fiji software (https://imagej.net/Fiji).

### Semiautomated sorting of area boundaries

Frequency domain boundaries for four tonotopic auditory cortical areas (A1, AAF, VAF, and A2) were determined semiautomatically for each tone frequency using trial-averaged intrinsic signal response maps, following a previously described approach (Narayanan et al., 2023). Response maps were first visualized with high intensity thresholds to facilitate segregation and identification of response centers for the four areas. Guided by these thresholded maps, coarse locations of frequency domains for each area (seed ROIs) were manually drawn. For each seed ROI, the domain centroid was defined as the mean of two locations: the point of peak response amplitude and the center of mass within the seed ROI. Next, around this domain centroid, the frequency domain mask was determined as the set of pixels surrounding each centroid whose signal intensity exceeded 60% of the peak amplitude. When frequency domain masks for the same stimulus overlapped between two cortical areas, a dividing line was drawn such that the distances from the dividing line to the two domain centroids were proportional to the peak response amplitudes of the respective domains. As an exception, overlap between the 3 kHz domains of A1 and VAF was permitted, reflecting the convergence of these areas at their low-frequency poles observed in most animals (Issa et al., 2014; Liu et al., 2019; Romero et al., 2020; Narayanan et al., 2023). The resulting area borders are largely independent of the initial placement of seed ROIs, as long as the seed ROIs include the peak response locations and are sufficiently separated from each other.

Once frequency domain masks were determined for three frequencies, these masks were combined to generate composite area masks for A1, AAF, VAF, and A2. After joining the binarized domain masks for the three frequencies, the boundaries were smoothed using opening and closing operations with disk-shaped elements of 30 and 150 pixels radius, respectively. In some animals, responses to one or two of the tested frequencies were not clearly detectable in specific cortical areas; in these cases, only the frequency domain masks for responsive stimuli were used. Finally, if the area masks for two cortical regions overlapped, a dividing line connecting the two intersection points was drawn. As above, overlap between A1 and VAF area masks was permitted.

### Calculation of relative variability in auditory cortex location

Reproducibility of tonotopic maps and harmonic response patterns shown in Fig. 1 was assessed by four experimenters who did not contribute to the imaging data in our previous studies (Kline et al., 2021, 2023), which reported preferential integration of multi-frequency sounds in A2. The dataset included mice of the following genotypes: B6-Cdh23 (*n* = 17), PV-Cre×B6-Cdh23 (*n* = 8), Ai9×Cdh23 (*n* = 3), VGAT-Cre×Cdh23 (*n* = 2), VGAT-Cre×Ai9 (*n* = 3), Sst-Cre×Ai9 (*n* = 1), PV-Cre (*n* = 3), VIP-Cre (*n* = 4), and PV-Flp × Calb2-Cre (*n* = 3). After generating area masks for A1, AAF, VAF, and A2 in each mouse, masks were rotated to align the skull ridge with the x-axis, resulting in approximate alignment of the x-axis with the anteroposterior stereotaxic axis (Narayanan et al., 2023). The auditory cortex center (AC center) was defined as the midpoint of the bounding box enclosing all identified auditory areas, and the positions of individual area masks were expressed relative to this reference. Probability distributions of individual cortical regions (Fig. 1D, middle panels) were visualized by overlaying area masks from all mice for each experimenter, applying a circular averaging filter (150 μm radius), and plotting the contour at 10% increments starting from 30%. Harmonic responses were mapped onto auditory cortical areas by applying the same rotation and recentering procedure to trial-averaged harmonic response maps for each animal and then averaging across all animals. For visualization (Fig. 1D, right panels), 50% contour lines of the probability distribution for each cortical region were overlaid onto the harmonic response heat maps.

The A2 localization index for harmonic responses in each mouse was calculated as

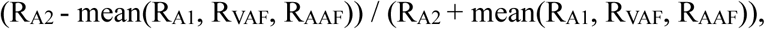

where R values represent harmonic response magnitudes in the corresponding cortical areas. This index ranges from +1 (exclusive localization to A2) to -1 (exclusive localization to non-A2 regions). For within-animal reproducibility analysis (Fig. 1F), only animals with at least eight trials per stimulus were included. Trials were split into two non-overlapping subsets by separating the trials with odd and even numbers, and area ROIs were independently determined for each subset (ROI 1 and ROI 2). The Overlap Index between these two sets of ROIs (Fig. 1F) was calculated as

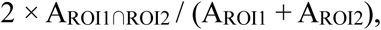

where A denotes the area size of individual ROIs. The index ranges from 0 (no overlap) to +1 (complete overlap). To account for small alignment errors, area ROIs were dilated by 50 μm prior to Overlap Index calculation. Area centroids were defined as the centers of mass of individual area masks, and absolute distances between centroid pairs from the two trial subsets were computed and averaged across the four cortical regions for each mouse.

### Quantification of changes in auditory cortical maps after window implantation

We reanalyzed mice from our previously published two-photon calcium imaging work together with newly collected animals, all of which were used for two-photon calcium imaging via virally expressed GCaMP. We additionally included mice whose data had been excluded from publication due to distorted cortical maps, insufficient GCaMP expression, or poor optical access caused by regrowth of thin connective tissue. The dataset included mice of the following genotypes: B6-Cdh23 (*n* = 4), Sst-Cre×B6-Cdh23 (*n* = 3), VGAT-Cre×Ai9×Cdh23 (*n* = 2), VGAT-Cre×Ai9 (*n* = 23), PV-Cre×Ai9 (*n* = 10), Sst-Cre×Ai9 (*n* = 4), Scnn1a-Cre×Ai9 (*n* = 2), and VGAT-Cre (*n* = 2). In all experiments, transcranial intrinsic signal imaging was performed before window implantation to guide viral injections and window placement. A second round of intrinsic signal imaging was conducted through the glass window approximately two weeks after window implantation, before proceeding with two-photon calcium imaging. Because intrinsic signal imaging in these animals was not originally intended for quantitative pre-post comparisons, some datasets exhibited insufficient signal-to-noise ratio for quantitative analysis due to technical factors such as limited trial numbers or bleeding from skull vasculature. These animals were excluded from the current analyses.

Area ROIs were determined independently from intrinsic signal maps acquired before and after window implantation. Minimal rotation of the fields of view between the two imaging sessions was assumed because the metal head bar was implanted prior to the first imaging session, and the same head-fixating apparatus was used for both sessions. Accordingly, we identified the optimal translation, without rotation, that maximized overlap between pre- and post-window area ROIs for each mouse. After translating pre-window ROIs into post-window coordinates, the Overlap Index and area centroid shifts were quantified as described above. Portions of pre-window ROIs that fell outside the cranial window in post-window coordinates were excluded before calculating the Overlap Index. Cranial window dimensions were measured by manually fitting an ellipse to images of the implanted window, and window area was calculated from the lengths of the major and minor axes.

To estimate the optimal ratio (r_opt_) between dorsoventral (D_DV_) and anteroposterior (D_AP_) window dimensions, we modeled the effective dimension governing compression in each animal as:

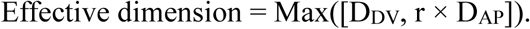

In this model, if D_DV_ exceeds r_opt_ × D_AP_, compression along the DV axis dominates; conversely, if r_opt_ × D_AP_ exceeds D_DV_, compression along the AP axis dominates. To identify r_opt_, the ratio r was varied from 0 to 2 at 0.01 steps, and the r value that maximized the coefficient of determination (R^2^) between the effective dimension and each map distortion measure (Overlap Index and area centroid shift distance) was determined, respectively.

The A2 preference index for the harmonics-to-tone response ratio was calculated for each mouse as:

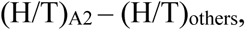

where H and T denote response magnitudes to harmonic stimuli and pure tones (mean of 3, 10, and 30 kHz tone responses), respectively, in individual areas. For analysis in Fig. 4B and 4D (A2 vs. others), (H/T) values were averaged across A1, AAF, and VAF. For other panels in Fig. 4D, the index was computed separately for pairwise comparisons between A2 and A1, AAF, or VAF.

The visualization of cortical geometry around the auditory cortex in the Allen CCF (Fig. 3A, Supplementary Fig. 3) used Brain Explorer 2 (https://mouse.brain-map.org/static/brainexplorer).

### Analysis of two-photon calcium imaging data

Lateral motion was corrected using cross-correlation-based image alignment. Regions of interest (ROIs) corresponding to individual cell bodies were detected automatically using Suite2P (Pachitariu et al., 2017) and supplemented by manual drawing. All ROIs were individually inspected and edited for appropriate shapes using a graphical user interface in Matlab. Two-photon imaging fields were aligned with the intrinsic signal imaging fields by comparing blood vessel patterns, and ROIs determined to be outside of the areal border determined by intrinsic imaging were excluded from analyses. Pixels within each ROI were averaged to derive a fluorescence time series, F_cell_measured_(t). To correct for background contamination, ring-shaped background ROIs (starting at 2 pixels and ending at 8 pixels from the border of the ROI) were created around each cell ROI. From this background ROI, pixels that contained cell bodies or processes from neighboring cells (detected as pixels showing large increases in dF/F uncorrelated to that of the cell ROI during the entire imaging session) were excluded. The remaining pixels were averaged to generate a background fluorescence time series, F_background_(t). The fluorescence signal for each cell was estimated as F(t) = F_cell_measured_(t) – 0.9 × F_background_(t). To ensure robust neuropil subtraction, only cell ROIs that were at least 3% brighter than their corresponding background ROIs were included in subsequent analyses. To compensate for slow drifts in baseline fluorescence intensity, a normalized trace was calculated as F_norm_(t) = F(t)/F_0_(t), where F_0_(t) represents a slowly varying baseline trace estimated by smoothing the inactive portions of F(t) using an iterative procedure.

Harmonic-evoked responses were quantified as the area under the curve of baseline-subtracted dF/F traces during a 1-s window following sound onset, and negative response amplitudes were set to zero. Cells were judged as significantly excited if they fulfilled both of the following two criteria: 1) dF/F had to exceed a fixed threshold value (3.3 times the standard deviation of the baseline period) consecutively for at least 0.5 s in more than half of the trials. 2) dF/F averaged across trials had to exceed a fixed threshold value consecutively for at least 0.5 s. Coincidence preference was quantified using mean dF/F traces averaged across at least five trials for each sound stimulus. The Coincidence Preference Index was calculated for each ROI-F_0_-Δonset combination only when significant excitatory responses were evoked by coincident and/or temporally shifted sounds as:

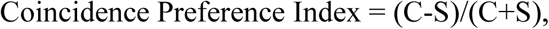

where C denotes the response amplitude to coincident sounds, and S denotes that to temporally shifted sounds. The indices for onset shifts of ±15 and ±30 ms were pooled across ROIs and compared between A1 and A2.

The responsiveness of individual neurons to pure tones was assessed using responses to 1-second pure tone stimuli. To facilitate direct comparison with a study reporting a low fraction of tone-responsive neurons (Liu et al., 2019), we first implemented their response-detection criterion as described. Specifically, for each trial, response magnitude was calculated as the dF/F value measured in a small window (±2 frames) centered on the maximum time point within 1 second after sound onset. For each neuron-stimulus combination, we estimated the 95% confidence interval of the median response magnitudes using bootstrapping (1000 resamples across trials). A neuron was classified as significantly excited if the lower bound of this confidence interval exceeded a threshold defined as 1.5 times the standard deviation during the baseline. When applied to our datasets, this criterion classified approximately 80% of cells as tone-excited, with a false positive rate in no-sound trials as high as 20%. To better control false positives, we therefore used a more conservative threshold of 2.5 times the standard deviation, which reduced the false positive rate to 5%. In addition, to reduce isolated threshold crossings by chance, we required that significant responses cluster within the frequency-intensity space; responses were accepted only if there were at least three significant responses within ±0.5 octaves (a total of 5 sound frequencies) and three sound levels. Using these stringent criteria, 38.1 ± 2.5% of A1 pyramidal neurons were responsive to at least one pure tone (Fig. 5G).

### Statistics and Reproducibility

Sample sizes were not predetermined by statistical methods. All key findings were replicated across multiple mice. Randomization or blinding were not performed, as experimental conditions were defined by surgical preparation and anatomical location rather than treatment assignment. Data are presented as individual data points or as mean ± SEM, as specified in the figure legends. Statistical differences between conditions were evaluated using two-sided non-parametric tests, specifically the Wilcoxon rank-sum test for unpaired data and the Wilcoxon signed-rank test for paired data. Bonferroni correction was applied for multiple comparisons, and corrected p-values were reported. Correlation coefficients and associated p-values were calculated as Pearson correlation using Matlab’s corrcoef function.

## Data availability

Source data for all figures will be provided as a supplementary data file attached to this paper.

## Code availability

This project did not develop broadly applicable software packages. However, code is available from the corresponding author upon request.

## Acknowledgements

This work was supported by the National Institute on Deafness and Other Communication Disorders (R01DC017516 to H.K.K.), the National Institute of Neurological Disorders and Stroke (R01NS128873 to H.K.K.; F31-NS111849 and T32-NS007431 to A.M.K), the Simons Foundation, the Eagles Autism Foundation, the Pew Biomedical Scholars Program, the Whitehall Foundation, the Klingenstein-Simons Fellowship (H.K.K.), and Foundation of Hope (H.K.K. and H.T.).

## Author Contributions

A.M.K. and H.K.K. designed the project. A.M.K. and H.T. performed calcium imaging. A.M.K., H.T., M.M., G.P.S., M.M.G, and H.K.K. performed intrinsic signal imaging. H.K.K. performed data analysis and wrote the manuscript, with inputs from all authors.

## Declaration of Interests

The authors declare no competing interests.

## Notes

### Competing Interest Statement

The authors have declared no competing interest.

