## Supplementary Figures for "Large cranial windows distort sensory maps and degrade feature integration in higher-order cortex"

**Supplementary Information**

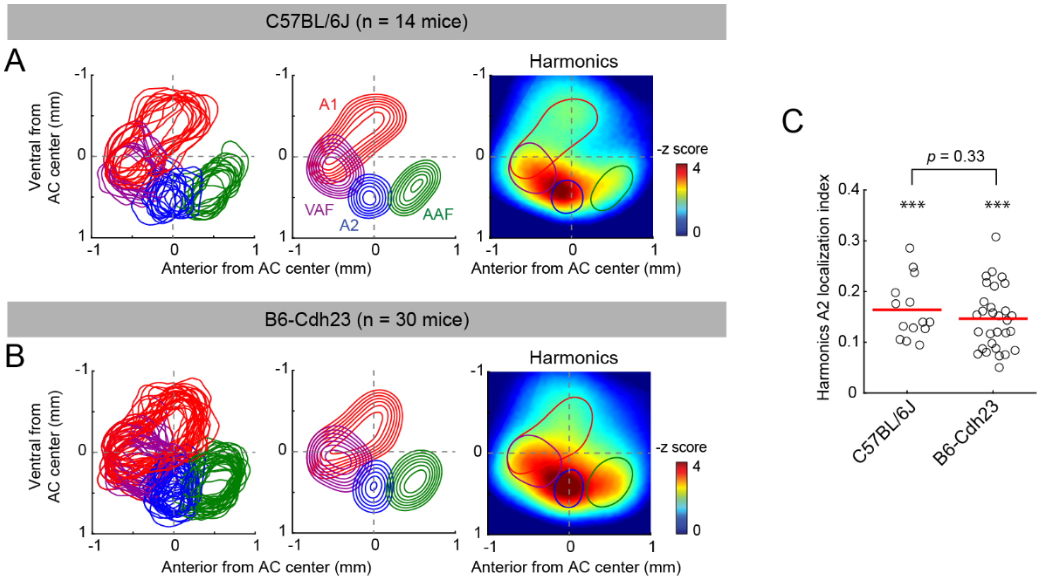


**Figure S1. Reproducible mapping of harmonic responses to A2 across mouse strains. (A)** Distributions of functionally identified area boundaries and harmonic responses obtained from mice on a C57BL/6J background (*n* = 14 mice). Left, cortical area boundaries from all mice superimposed in coordinates around the AC center. Middle, probability distributions of area masks for individual cortical regions. Contours indicate 10% steps, starting from 30%. Right, heat map of harmonic responses averaged across animals, overlaid with the 50% contour of the area probability distribution. **(B)** Same as in (A), but for mice on a B6-Cdh23 background, which do not exhibit age-related hearing loss (*n* = 30 mice). **(C)** Scatter plot showing the A2-localization index of harmonic responses for individual strains. Red lines indicate the mean. ***$p < 0.001$ (two-sided Wilcoxon signed-rank test with Bonferroni correction). C57BL/6J vs. B6-Cdh23: $p = 0.33$ (two-sided Wilcoxon rank-sum test).

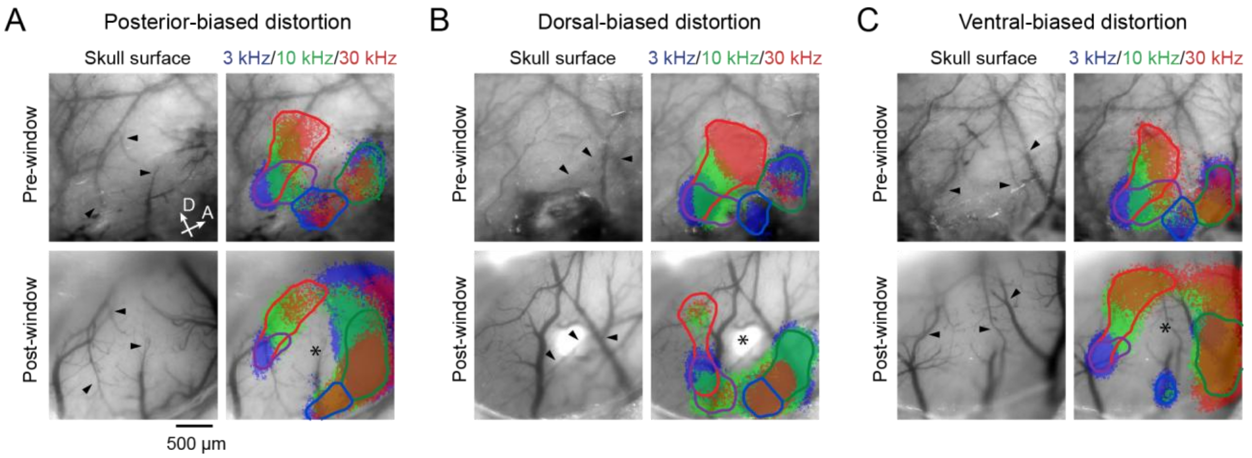


**Figure S2. Example mice exhibiting side-biased distortion after cranial window implantation. (A–C)** Left, images of cortical vasculature acquired before (top) and after (bottom) cranial window implantation in the same mouse. Black arrowheads indicate the same blood vessels identified across imaging sessions. Right, thresholded intrinsic signal responses to pure tones at 3, 10, and 30 kHz acquired before (top) and after (bottom) cranial window implantation, with overlaid autosorted area boundaries. Asterisks denote central regions with reduced sound responses following window implantation. **(A)** Example mouse showing map distortion biased toward the posterior side. **(B)** Example mouse showing map distortion biased toward the dorsal side. **(C)** Example mouse showing map distortion biased toward the ventral side.

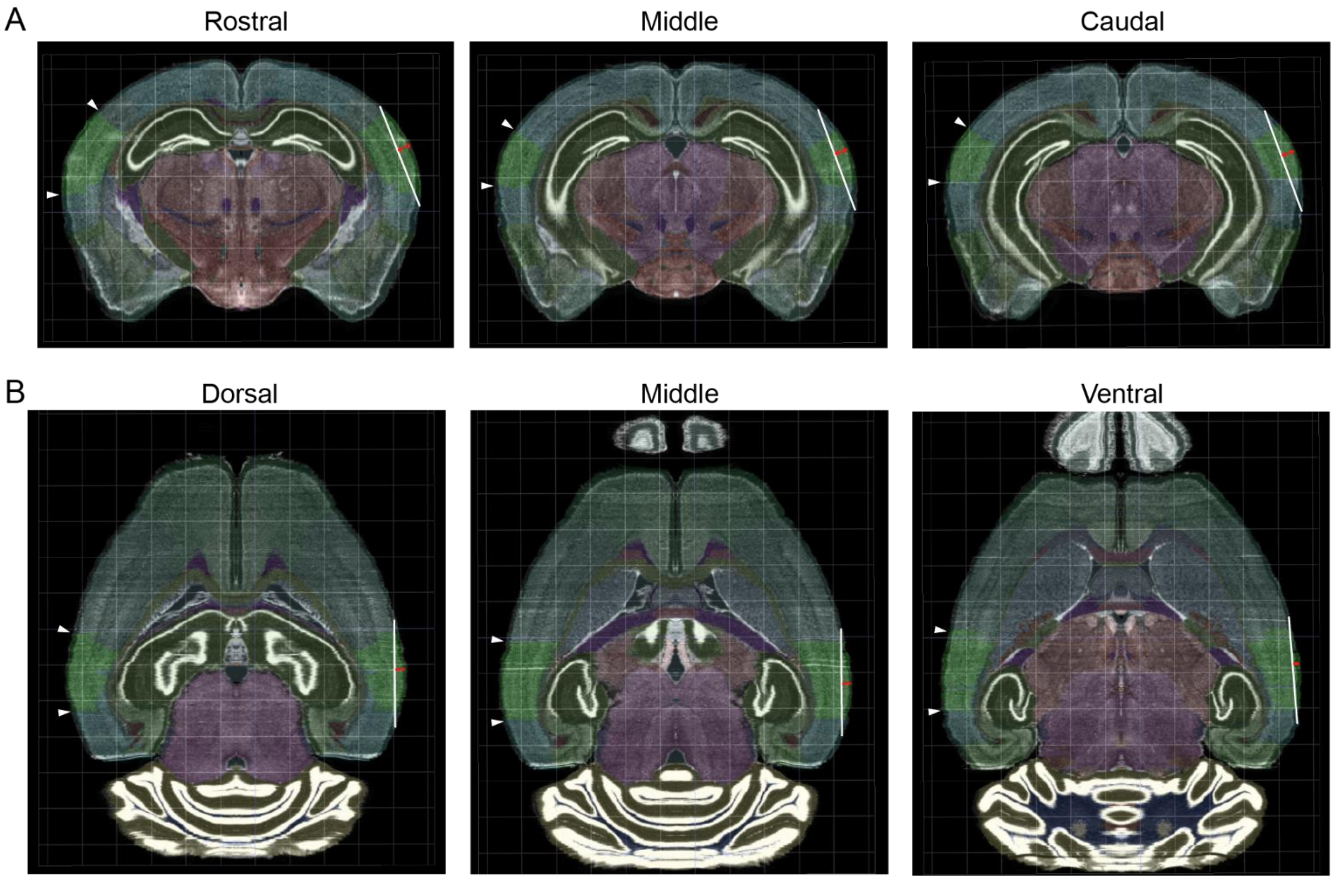


**Figure S3. Predicted cortical compression by a 3-mm cranial window over the auditory cortex. (A)** Coronal sections from the Allen Common Coordinate Framework (CCF) containing rostral, middle, and caudal portions of the auditory cortex. White lines indicate the diameter of a 3-mm cranial window positioned against the curved cortical surface. Red arrows indicate the resulting compression. White arrowheads on the contralateral hemisphere and green shading denote the boundaries of the auditory cortex as defined in the Allen CCF. **(B)** Same as in (A), but for horizontal sections containing dorsal, middle, and ventral portions of the auditory cortex. Across positions, cortical compression along the dorsoventral axis is predicted to dominate over compression along the anteroposterior axis.

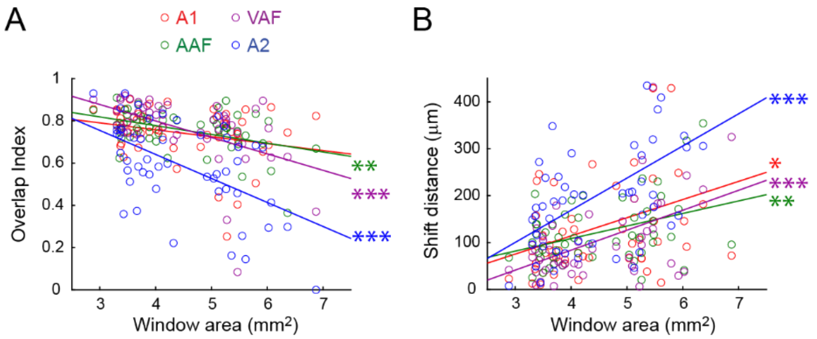


**Figure S4. Window size-dependence of distortions in individual auditory cortical areas. (A)** Overlap Index between pre- and post-window area ROIs calculated separately for individual auditory cortical regions, plotted as a function of cranial window area for individual mice. Colored lines indicate linear regression fits. $n$ = 50 mice. A1: $R$ = -0.258, $p$ = 0.0704; AAF: $R$ = -0.421, **$p$ = 0.0023; VAF: $R$ = -0.506, ***$p$ = 0.00018; A2: $R$ = -0.538, ***$p$ = $5.60 \times 10^{-5}$. **(B)** Centroid shift distances between pre- and post-window area ROIs for individual auditory cortical regions, plotted as a function of cranial window area for individual mice. $n$ = 50 mice. A1: $R$ = 0.326, *$p$ = 0.0208; AAF: $R$ = 0.387, **$p$ = 0.0055; VAF: $R$ = 0.501, ***$p$ = 0.00021; A2: $R$ = 0.530, ***$p$ = $7.69 \times 10^{-5}$.

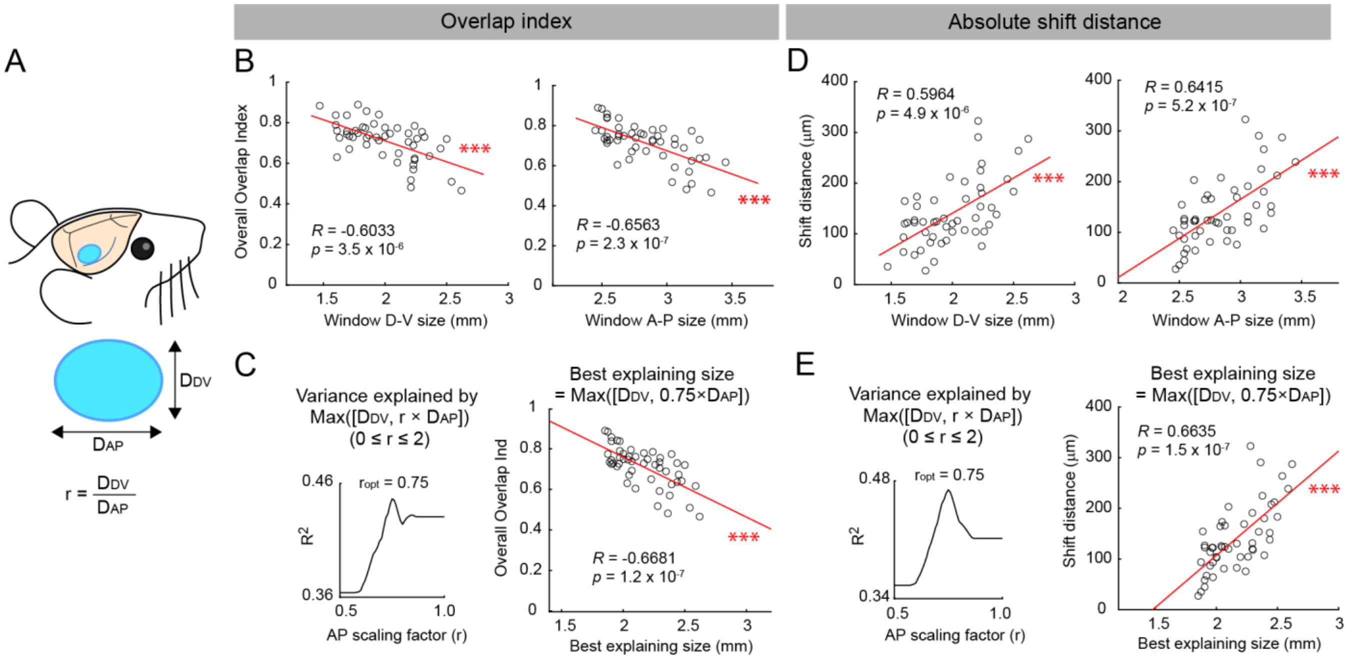


**Figure S5. Determination of the optimal ratio between dorsoventral and anteroposterior cranial window dimensions. (A)** Schematic illustrating an oval cranial window elongated along the anteroposterior axis, which is predicted to minimize cortical compression based on the geometry of the Allen CCF. The parameter r denotes the ratio between the dorsoventral ($D_{DV}$) and anteroposterior ($D_{AP}$) window dimensions. **(B)** Overall Overlap Index between pre- and post-window area ROIs plotted as a function of $D_{DV}$ (left) and $D_{AP}$ (right). Red lines indicate linear regression fits. $n = 50$ mice. $D_{DV}$: $R = -0.603$, ***$p = 3.5 \times 10^{-6}$; $D_{AP}$: $R = -0.656$, ***$p = 2.3 \times 10^{-7}$. **(C)** Left, coefficient of determination ($R^2$) between the Overall Overlap Index and the effective window dimension plotted as a function of r. The optimal ratio ($r_{opt}$) was identified as 0.75. Right, Overall Overlap Index plotted as a function of the best explaining effective window size. $R = -0.668$, ***$p = 1.2 \times 10^{-7}$. **(D, E)** Same as in (B, C), but for absolute centroid shift distances between pre- and post-window area ROIs. **(D)** Centroid shift distance plotted as a function of $D_{DV}$ (left) and $D_{AP}$ (right). $n = 50$ mice. $D_{DV}$: $R = 0.596$, ***$p = 4.9 \times 10^{-6}$; $D_{AP}$: $R = 0.642$, ***$p = 5.2 \times 10^{-7}$. **(E)** $R^2$ between centroid shift distance and effective window dimension plotted as a function of r, identifying $r_{opt}$ as 0.75. Right, Centroid shift distance plotted as a function of the best-explaining effective window size. $R = 0.664$, ***$p = 1.5 \times 10^{-7}$.
